# Exploring adaptive introgression in modern human circadian rhythm genes

**DOI:** 10.1101/2024.08.12.607550

**Authors:** Christopher Kendall, Amin Nooranikhojasteh, Guilherme Debortoli, Vinicius Cauê Furlan Roberto, Marla Mendes, David Samson, Esteban Parra, Bence Viola, Michael A. Schillaci

**Author notes:** Corresponding author (CK).

## Abstract

Interbreeding between modern humans and archaic hominins, including Neanderthals and Denisovans, occurred as modern humans migrated outside of Africa. Here, we report on evidence of introgression from archaic hominins within genomic regions associated with circadian rhythm and chronotype using 76 worldwide modern human populations from the Human Genome Diversity Project and 1000 Genomes Project. We calculated the extent of regions indicative of adaptive introgression across the autosomes and identified regions that are suggested to be under positive selection. We tested for evidence of a latitudinal cline within 36 core haplotypes along with presenting the likely archaic donor for each of these haplotypes. We identified 265 independent segments that overlap genes described as having a circadian rhythm component or contain variants and segments previously identified as being associated with circadian rhythm or chronotype. Within these segments we found 1,729 archaically derived variants with allele frequencies of at least 40% intersecting 303 genes and intergenic segments. Seventeen of these segments show evidence of positive selection, three of which are found within our core haplotypes. We found that many of our genes are associated with the immune system or gastrointestinal function. Additionally, variants associated with complex traits such as schizophrenia and bipolar disorder are present within our adaptively introgressed regions. Lastly, genes and markers associated with sleep and chronotype phenotypes and serotonin pathways were also found in our adaptive introgression results, potentially signalling selection on genes related to seasonal light variation as modern humans migrated into new environments after leaving Africa.

**Author summary:** As modern humans migrated out of Africa, they encountered archaic hominins, the Neanderthals and Denisovans, and interbred with them. Signatures of these admixture events can be found in populations across the world. The result of these admixture events has shaped modern human evolution regarding high altitude adaptation, immune function, and skin and hair colour, to name a few. However, much of this information has been gathered with a focus on Eurasian populations using the 1000 Genomes Project samples. Here, we take advantage of newly published resources from 76 worldwide modern human populations to investigate how strongly a role admixture played on modern human circadian rhythm genes. Circadian rhythms have been tied to sleep-wake regulation, immune function, and digestive health. We find evidence for adaptive introgression in over 300 genes and intergenic segments. Many of these genes, like *AMIGO2*, are associated with complex traits such as schizophrenia and bipolar disorder or with immune system function, like *JAK1*. Some of these traits have been previously described before regarding archaic admixture. Interestingly, many of these associated traits are influenced by circadian rhythm oscillations, providing a new perspective on interpreting these findings.

## Introduction

A little over a decade ago, human evolutionary history was completely reshaped by the discovery that anatomically modern humans interbred with our archaic cousins, the Neanderthals [1], and their enigmatic sister species, the Denisovans [2]. These were not isolated events, and evidence has been uncovered that there were several admixture events occurring sporadically over thousands of years and across diverse geographic areas [3–9]. Signatures of these events are left in our genome with estimates that every non-African alive today has on average just below 2% of their DNA shared with Neanderthals [10]. Denisovan signatures in modern humans are generally lower, on average below 1% [11–12]. However, some Oceanic populations have been noted to have nearly 5% of their DNA composed of Denisovan-introgressed regions [2, 8, 13].

A number of these archaic introgressed regions are believed to have been adaptive and have been brought to elevated frequencies in modern human populations. Several of the most notable examples are variants within the *EPAS1* gene in Tibetan populations that confer adaptation to high altitude environments derived from Denisovans [14], immunity and HLA-controlling regions likely giving rise to disease resistance to modern humans as they expanded into new territories after leaving Africa [15–18], and the various skin, hair, and keratin linked regions introgressed from Neanderthals that have been highlighted in a number of studies [4, 19–21]. While these are often referenced, there are many other introgressed loci that have been associated with traits such as type 2 diabetes risk [22], height, likelihood of being a smoker, mood [20], and mental health [23], to name a few.

Recently, the discovery of introgressed variants within genes involved in circadian rhythm and chronotype expression has become a new area of focus [20–21, 24–26]. Single nucleotide polymorphisms (SNPs) identified to be associated with chronotype, daytime napping, narcolepsy, lethargy, willingness to get up in the morning, sleep duration, and insomnia have been suggested to be the product of introgression from Neanderthals [20]. In a meta-analysis using previously published genome wide association study (GWAS) data identifying archaic introgressed loci, modern human genetic variants associated with being a morning person were shown to be shared with the Altai Neanderthal [21]. Further, analysis of GWAS data from several biobanks found positive association with archaically identified SNPs related to sleep traits such as narcolepsy, daytime napping, sleep duration, willingness to get up in the morning, and chronotype [25]. In a recent analysis using a combination of previously published variants identified as being archaically-introgressed into modern humans, it was found that modern humans and archaic hominins differed in their circadian rhythm genes, including alternative splicing events and regulatory divergence [26]. Additionally, the authors noted that over 3,800 variants were associated with regulation on circadian genes, these variants are more likely to be expression quantitative trait loci (eQTL) for circadian rhythm genes than by chance, and that 47 circadian genes show evidence of adaptive introgression [26]. Lastly, the authors of that study state that introgressed variants are associated with having a morningness chronotype and that some introgressed variants are distributed across a latitudinal cline [26].

The circadian rhythm is the cyclic oscillator of a 24-hour period, which has remained relatively conserved across most of the animal kingdom [27] and has been proposed to be a core controller of sleep and wake cycles [28–32]. To do this, the suprachiasmatic nucleus, located in the hypothalamus, uses external light stimuli to reset itself along a day-night cycle [31]. Output from the suprachiasmatic nucleus goes to the ventral subparaventricular zone and regulates this information into daily cycles of wakefulness and sleep [27, 31], that then falls across the natural 24-hour circadian rhythm. In addition to controlling sleep and wake cycles, several review articles have highlighted the link between circadian rhythm and gastrointestinal processes [33–34] and immune function [35–37].

The preference for how late someone stays awake is influenced by chronotype, that is, morning people who tend to go to bed and rise earlier, and those who show an evening preference and go to bed and rise later [32, 38]. Three different GWAS datasets have independently identified four genes that support associations with chronotype, *PER2*, *RGS16*, *FBXL13*, and *AK5* [32]. Additional studies increased the number of chronotype-linked variants which were associated with previously unidentified genes, such as *PER1*, *CRY1*, and *ARNTL* [39]. Links between daytime light exposure and chronotype expression in modern humans have also been suggested on candidate gene *ARL14EP* [40].

For this study we used the gnomAD 1000 Genomes Project (1KGP) and Human Genome Diversity Project (HGDP) phased callset [41] to identify regions of archaic introgression from an expanded worldwide population dataset (n=76 populations). Specifically, we used SPrime [7] to identify genomic regions in these populations with an interest in markers associated with circadian rhythm and chronotype phenotypic expression that are the result of introgression from archaic hominins into modern humans. To explore if any of these regions would contain signatures of adaptive introgression on genes showing circadian rhythm patterns, we extracted the modern human genes from the Circadian Genome Database (CGDB) [42]. We also included in our analysis data from 4 previous studies that discussed chronotype and archaic introgression, and lastly, included regions from the NHGRI-EBI GWAS Catalogue [43] that were associated with circadian rhythm, chronotype, and sleep to identify regions of adaptive introgression associated with these phenotypes found in our dataset. We highlight more than 1,700 variants at elevated frequencies in our dataset that fall within 265 independent, genome-wide windows in non-African populations. Our research hypothesised that like prior studies [26], circadian rhythm and/or chronotype loci at levels suggestive of adaptive introgression will have phenotypic expressions largely driven by latitude, which we investigated using genome wide data. We utilised the four high coverage archaic genomes from the Altai Denisovan [2], Altai Neanderthal [3], Vindija Neanderthal [44], and the Chagyrskaya Neanderthal [45] for our analyses to determine if sample-specific signatures could be identified and in an attempt to gauge which archaic group was the likely donor population to core haplotypes found in our data.

Prior data has illustrated latitudinal clines with a genetic basis [46–47] and some of these are also detected within archaically derived segments [24, 26]. For this study, we predict that the frequency of circadian rhythm and chronotype-related archaic variants will vary with latitude, with higher latitudes experiencing more extreme seasonal variations in daylight exhibiting a greater frequency of introgressed variants associated with morningness chronotype and enhanced light sensitivity. Given the geographic distribution of Neanderthals and Denisovans [2, 48–50] we further hypothesise that archaic hominins, who inhabited higher latitudes, contributed a greater number of circadian rhythm and chronotype-related genetic variants to modern humans as humans left Africa. This prediction is grounded in the need for adaptations to significant seasonal light variations in these regions. Additionally, we predict that these archaic groups will be associated with a higher prevalence of variants that enhance serotonin synthesis, such as within the *DDC* gene [51], supporting adaptations to extreme seasonal variations in light. Variants associated with higher serotonin synthesis will likely correlate with phenotypic traits such as increased wakefulness during daylight hours and potentially reduced REM sleep. Accordingly, these variants are predicted to be more pronounced in these populations as this adaptation might help mitigate seasonal affective disorder (SAD) and maintain stable circadian rhythms despite fluctuating daylight hours.

## Results

### Introgression

Our analysis was able to recover a large number of variants reported in prior studies, showing the effectiveness of our pipeline (S1 Text). From the Dannemann and Kelso (2017) paper [20] reporting archaic SNPs found in modern humans, 1,787 variants were recovered in our dataset, while from Dannemann et al. (2022) [25], 1,415 SNPs were identified as archaically-introgressed in our dataset. We were able to extract 4,255 variants from the genomic windows associated with archaic introgression reported in McArthur et al. (2021) [21]. Lastly, from the data reported in Velazquez-Arcelay et al. (2023) [26] providing archaic-derived circadian and chronotype loci, we were able to recover 9,605 variants in our dataset. In short, our pipeline was successful at recovering previously reported instances of introgression associated with circadian rhythm and chronotype. We were also able to recover previously documented introgression patterns, including both the Chagyrskaya and Vindija Neanderthals being more closely related to the introgressed Neanderthal DNA in modern humans [44–45], and higher levels of Denisovan ancestry in Oceanic populations relative to other modern human groups [2] (Fig 1). Next, we recovered 64,834 putative archaic variants overlapping the genes listed in the CGDB [42]. Our analysis of the GWAS data [43] for chronotype and sleep-associated traits yielded a very small number of archaically derived hits, with only 71 markers found in our results.

**Fig 1.**
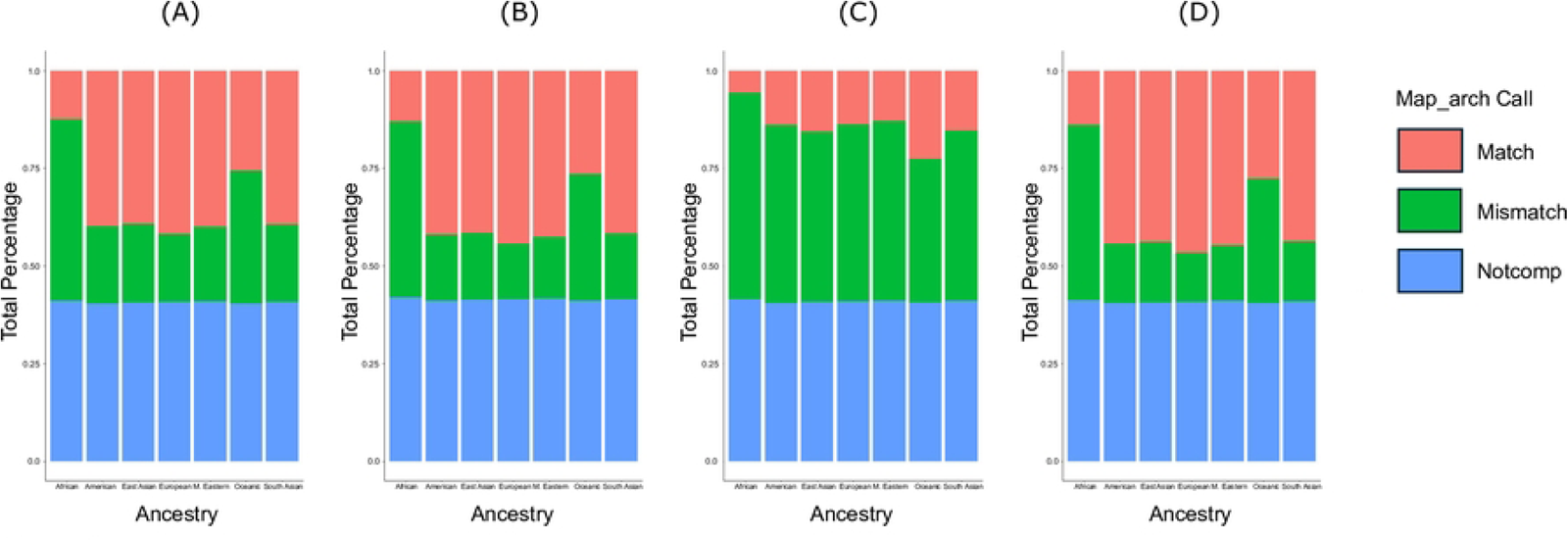
Archaic variant recovery. Putative archaic variants recovered in our pipeline. Colours denote map_arch labelling [52] and are representing the percentage of total recovered variants for the (A) Altai Neanderthal, (B) Chagyrskaya Neanderthal, (C) Vindija Neanderthal, and (D) Denisovan. The plot was generated using the ggplot2 package [53] package in R v4.1.2 [54].

We were first interested in the identification of regions of the modern human genome associated with circadian rhythm and chronotype with atypically high archaic allele frequencies. We identified 265 independent, non-overlapping segments in our 62 non-African populations (S1 Table), where each segment overlaps at least one variant or window associated with circadian rhythm or chronotype, or overlaps the genes listed in the CGDB [42], and the archaic variant has an allele frequency ≥ 40% in at least one population within our analysis. Within the Americas, there are 127 segments that pass these criteria, 123 in East Asia, 44 in Europe, 12 in the Middle East, 88 in Oceania, and 16 in South Asia (S1 Table). Within these segments, we noted 1,094 independent variants combined between all populations. When we filtered these regions to include only variants that are the maximum archaic allele frequency in their segment, we found there are 209 genes and intergenic regions within 131 independent segments (S2 Table). S2 Table also includes the population intersection results and gene information for the circadian rhythm or chronotype associated variant per population-specific segment. In S3 Table we expand upon the contents of S2 Table and include all variants with archaic allele frequencies ≥ 40%, including those that may not be the maximum archaic allele in their respective segment. S3 Table highlights a total of 1,729 variants that are found within 303 genes and intergenic regions.

### Core haplotypes and evidence of positive selection

After generating our core haplotypes (Materials and Methods), we were left with 36 regions of interest for further exploration (S4 Table). Twenty-four (∼67%) of these regions fall within genes while the remaining 12 (∼33%) are intergenic (S4 Table). After running RAiSD [55], 17 of these regions show evidence of positive selection (S4 Table). However, only three of these regions, *CCR9* in the Indian Telugu in the U.K. (ITU) population, the larger *CEACAM1-LIPE-AS1* cluster in the Papuans, and *JAK1* in the Melanesians showed signatures of positive selection within the core haplotype. The remaining 14 haplotypes had positive selection signatures that fell within the introgressed population-specific segment, but outside of the core haplotype (S4 Table). After filtering for haplotypes of interest (see S1 Text), *CCR9* has 59 haplotypes, *CEACAM1-LIPE-AS1* has 63 haplotypes, and *JAK1* has 68 haplotypes in their core regions, respectively. Haplotype networks and ancestral recombination graphs (see S1 Text) for *CCR9, CEACAM1-LIPE-AS1*, and *JAK1* are shown in Figs 2 and 3. The output window from RAiSD [55] where positive selection was detected can be seen in S4 Table for core haplotypes where selection exists, while the contour plots [52] for the three main core haplotypes are shown in Fig 4. Additionally, we identified the most probable archaic donor for each of the 36 core haplotypes, which is provided in S4 Table.

**Fig 2.**
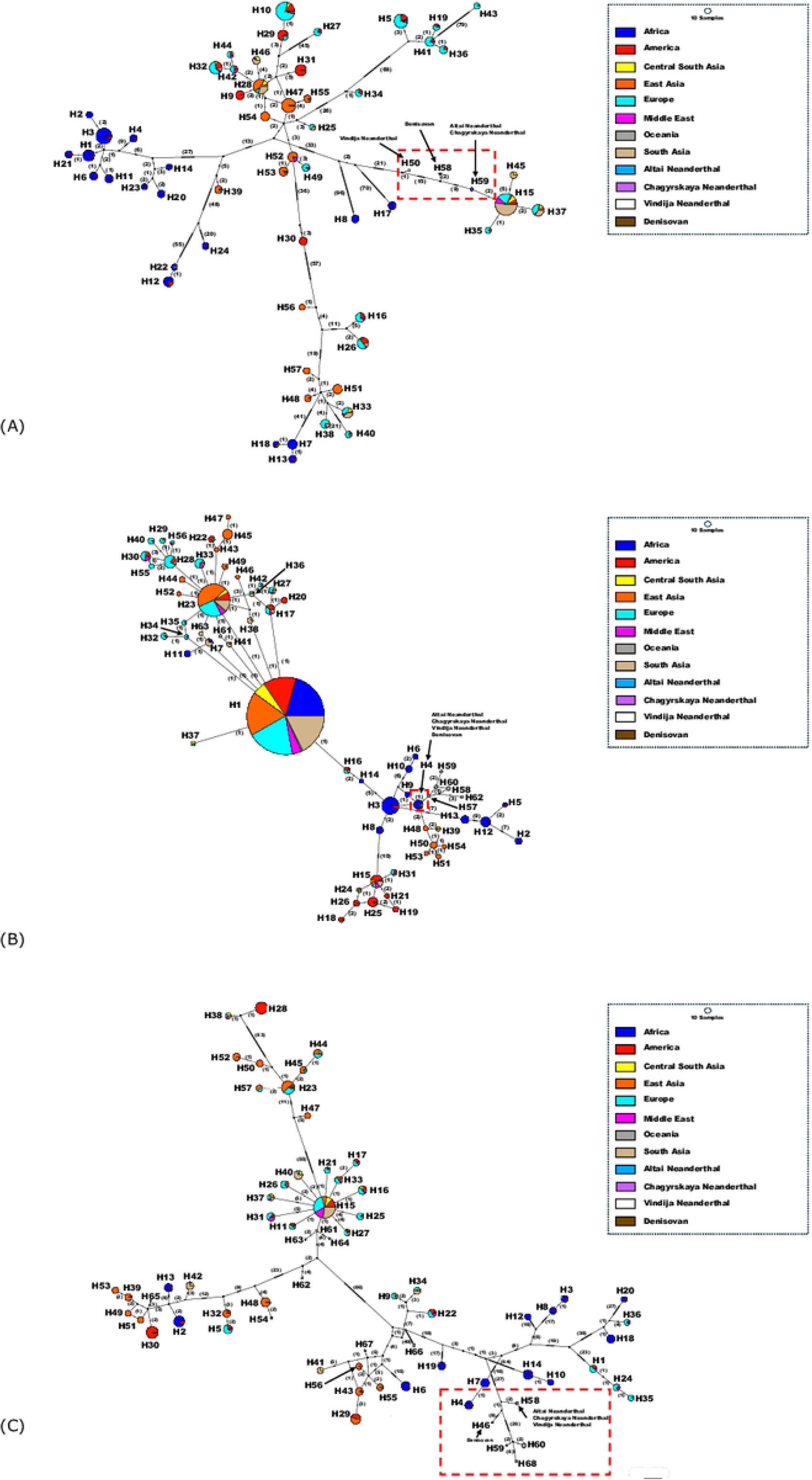
Haplotype networks of main core haplotypes. Haplotype networks for (A) *CCR9*, (B) *CEACAM1-LIPE-AS1*, and (C) *JAK1*, our three core haplotypes with evidence of positive selection within their core region. Red boxes highlight the location of archaic haplotypes. The number of mutations along each edge between nodes is shown in brackets. Plots were generated using PopArt v1.7 [56].

**Fig 3.**
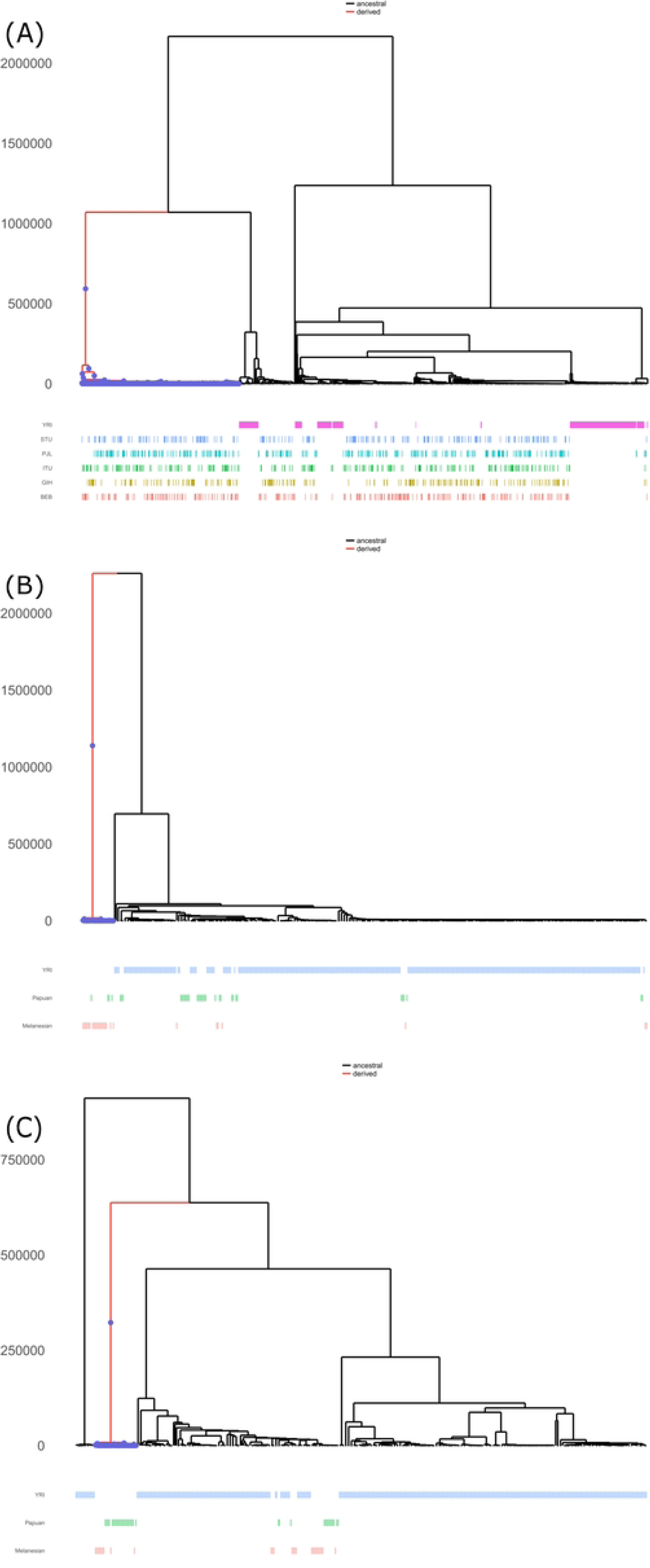
Ancestral recombination graphs of main core haplotypes. Ancestral recombination graphs for variants within core haplotypes with evidence of positive selection that are matches to the archaic allele and show patterns associated with archaic ancestry. These patterns include origins at least 1,000,000 years ago and long, non-recombining branches with recent expansion within modern human populations associated with derived mutations. (A) rs71327015 inside of the *CCR9* core haplotype in South Asian populations and rs377425962 inside of the *JAK1* core haplotype within Oceanic populations, respectively. rs184528844 inside of the *CEACAM1-LIPE-AS1* core haplotype in Oceanic groups, which shows an archaic-like branching event despite emergence after the split of archaic hominins and modern humans. Analysis and graphs were generated using Relate v1.2.1 [57].

**Fig 4.**
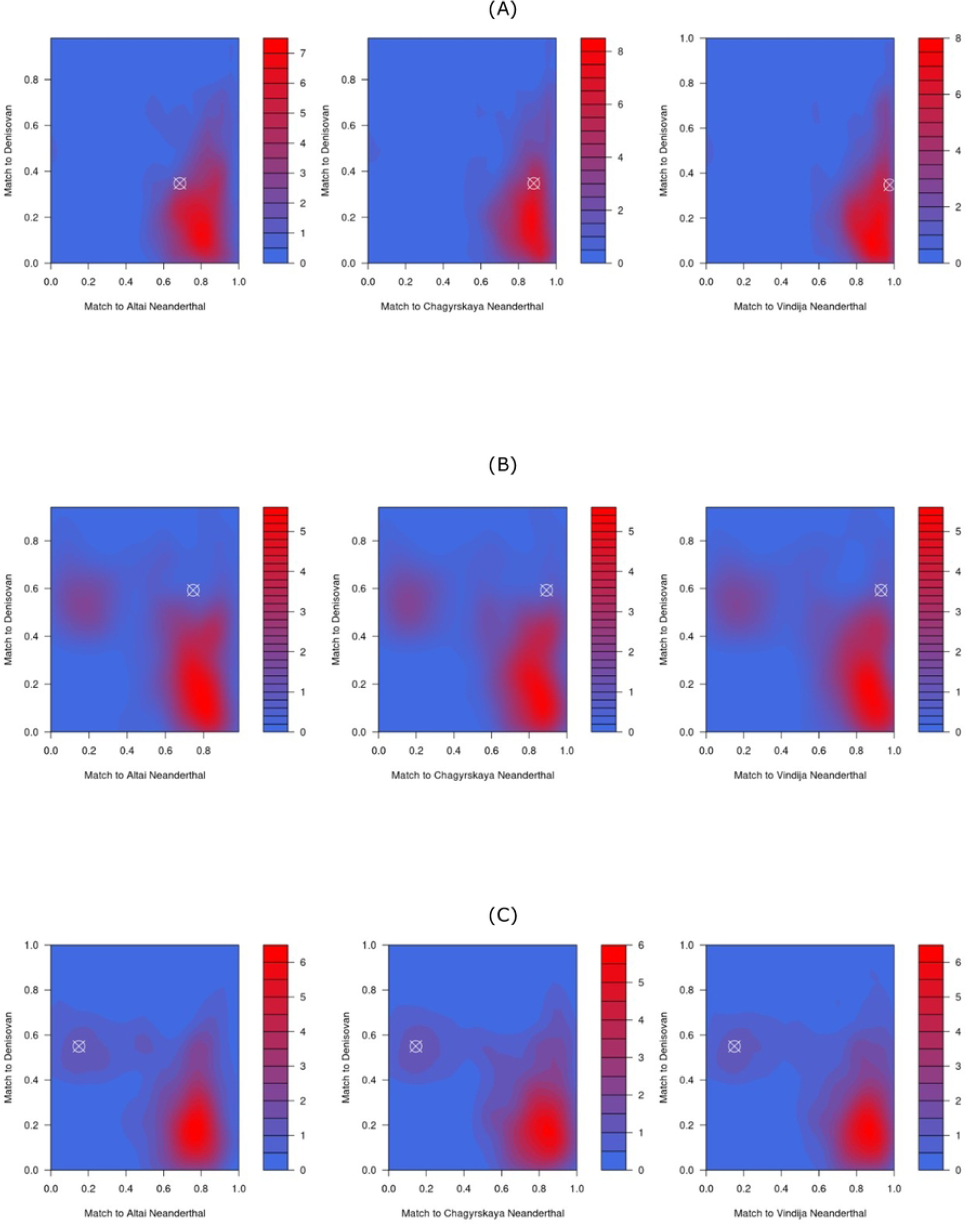
Archaic donor population of main core haplotypes. Contour plots based on the match/mismatch ratios of each putative archaic segment genome wide after filtering for authentic segments. The location of the segment containing the gene is identified by a white crosshair. Heatmap is coloured by the density of segments from both the Neanderthal and Denisovan samples at each match/mismatch ratio, with the archaic donor population being the archaic sample with the highest match/mismatch ratio. (A) *CCR9*’s segment is most like the Vindija Neanderthal, (B) *CEACAM1-LIPE-AS1* is of Neanderthal affinity generally, and (C) *JAK1* is most similar to the Denisovan. Plots were based upon the scripts provided by Zhou and Browning (2021) [52] using the kde2d function from the MASS library [58] in R [54].

### SNP annotations

We wanted to explore in more detail potential associations of the archaic variants provided in S3 Table to better understand the implications of our results. To do this we used the NHGRI-EBI GWAS Catalogue [43] and downloaded the association table results and matched the target SNPs within our identified regions against those found in the catalogue. In total, we identified 37 putative archaic SNPs that were found at frequencies ≥ 40% in our results and had genome-wide significant p-values in the GWAS catalogue. These variants are provided in S5 Table. We expanded our annotation analysis to include variant matches that may not reach genome-wide significant thresholds by exploring variants in our data that matched results found in SNPnexus [59–60] and the Variant Effect Predictor (VEP) [61]. These results are included in S6 Table.

## Discussion

### No evidence of latitude cline within core haplotypes

Several prior studies have provided evidence that circadian rhythm or chronotype associated genes and variants will often exhibit a latitudinal cline. Specifically, phenotypes related to chronotype based on latitude have been identified in modern human populations [46–47] and have also been described in regions introgressed from archaic populations [24, 26]. The link between latitude and circadian rhythm has been established in plants, animals, and insects to varying degrees [62–64]. Considering these claims, we tested if any evidence of a latitudinal cline could be found in our core haplotypes that displayed evidence of positive selection (S4 Table). We extracted the maximum archaic allele frequency from each population-specific segment intersecting our core haplotypes and plotted them based on longitude and latitude. Interestingly, and contrary to our prediction, we found no clear evidence of a latitude cline (Fig 5; S7 Table; S1-30 Figs). Four of these core haplotypes (*CEACAM1-LIPE-AS1*, *LINC01107-LINC01937*, *ROR2*, *TLR1*) have significant p-values (p ≤ 0.05) when testing for the relationship between maximum archaic allele frequency and latitude, but the relationship is not very strong with r^2^ values of 0.063541, 0.058346, 0.067914, and 0.23862, respectively (S7 Table). Most of the relationships are also counter to our hypothesis, where higher latitude groups, such as the British from England and Scotland (GBR) and Finnish in Finland (FIN) have lower allele frequencies than many populations from middle and low latitude regions (S7 Table). Only six core haplotypes (*CCR9*, *ENSG00000286749*, *LINC01107-LINC01937*, *RN7SL423P-ENSG00000232337*, *ROR2*, and *TLR1*) show positive relationships between maximum archaic allele frequencies and latitude (S2, S4, S8, S10-S11, S15 Figs), however, the *CCR9*, *ENSG00000286749*, *LINC01107-LINC01937*, and *RN7SL423P-ENSG00000232337* relationships are not significant (S7 Table). Further, our haplotype networks and ancestral recombination graphs also support this with no clear latitude signature displayed for our core haplotypes with evidence of positive selection within their core region (Figs 2-3; S31, S33, S35 Figs).

Despite a lack of a clear latitudinal pattern, we found evidence of geographic grouping of some core haplotypes. For instance, the *CEACAM1-LIPE-AS1* and *JAK1* frequency maps show a clear bias towards Oceania, with allele frequencies being greater than 40%, while these frequencies are nearly, or entirely, absent in the rest of our populations (Figs 5A-5B). Similar patterning can be seen in the *AMIGO2* frequency map, where elevated allele frequencies are isolated to Asia and Oceania (Fig 5C). We document high frequencies within Oceanic and South Asian populations in the *CCR9* frequency map, which also has some moderately elevated allele frequencies within Europe and some parts of the Americas (Fig 5D), and a South Asia to Europe band of high-frequency alleles in *RN7SL423P-ENSG00000232337* (Fig 5E). Some of these patterns are made very clear in our haplotype networks, where, for example in the *CCR9* network, geographic pockets of haplotypes are noted (Fig 2A). Within this haplotype network, most of the African haplotypes are separated from the other haplotypes, indicative of separate evolutionary histories within these putatively introgressed, high frequency regions. Much of the clustering is based on previously described levels of allele sharing between archaic populations and modern humans. In the *CEACAM1-LIPE-AS1* network (Fig 2B), contrasting to *CCR9*, the archaic haplotype shares an African haplotype, and is only one mutation away from two other African haplotypes. This may suggest that the archaic haplotype in this segment was derived from modern humans first, a recently discussed hypothesis in other genomic regions [10]. We would not expect to see many of these described groupings if circadian patterns related to latitude were solely controlling expression of these phenotypes. Lastly, our ancestral recombination graphs also support our inference, with edges and nodes shared between populations with strongly different latitudes (S31, S33, S35 Figs).

**Fig 5.**
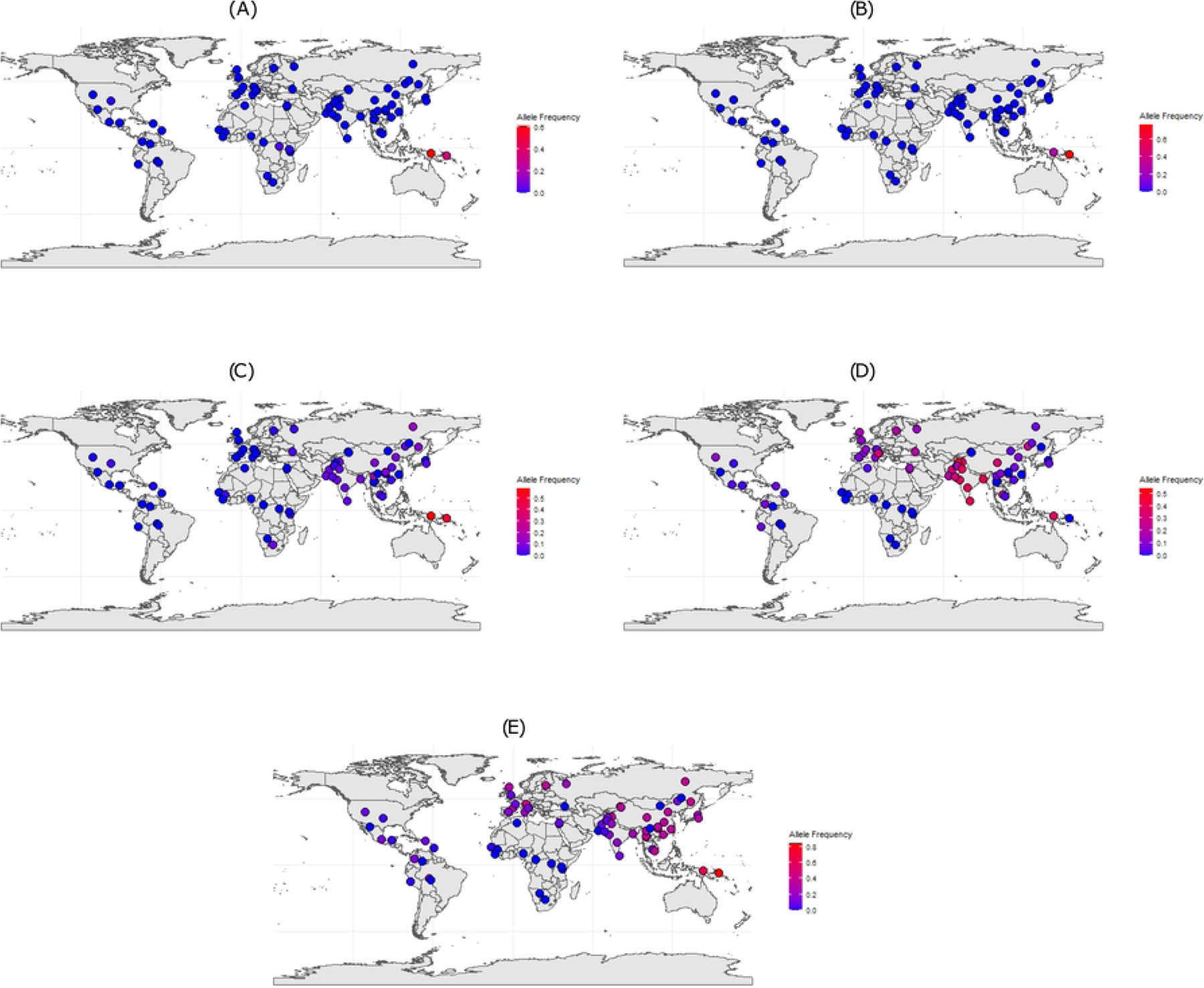
Frequency maps of core haplotypes with evidence of positive selection. Frequency distribution maps generated by extracting the maximum archaic allele frequency from each core haplotype that had evidence of positive selection generally within its putatively derived segment. Here we display a subset of these with interesting geographic patterns. Both (A) *CEACAM1-LIPE-AS1* and (B) *JAK1* are nearly exclusively found in Oceanic populations. (C) *AMIGO2* and (D) *CCR9* exhibit higher frequencies within Asia and Oceania. (E) *RN7SL423P-ENSG00000232337* shows a band across South Asia into Europe of high-frequency variants. Our analysis displays no clear latitudinal cline within our core haplotypes with evidence of positive selection. Plots were generated with the rnaturalearth [65], sf [66–67], and ggplot2 [53] packages in R [54].

### Serotonin associated genes with evidence of adaptive introgression

We found mixed results supporting the hypothesis that archaic populations contributed serotonin-associated variants to modern humans due to differing exposure to seasonal light variation, and further, that these variants will likely have higher instances of chronotype, mood, and sleep associated phenotypes. We downloaded from GeneCards [68] 375 genes that were associated with serotonin and intersected these against our list of 303 genes overlapping variants with archaic allele frequencies ≥ 40% (S3 Table). Four genes and intergenic regions, *CHST11*, *ENSG00000276064-HTR1B*, *MECOM*, and *TBC1D1* that have been previously linked with serotonin also have signatures of adaptive introgression in modern humans within our dataset (S3 Table). One of these genes, *CHST11*, is one of our core haplotypes without evidence of positive selection (S4 Table). Within these genes, there is no signature found regarding chronotype or sleep phenotypes according to our annotation results (S8 Table). However, *CHST11* has been associated with schizophrenia and bipolar disorder [69], celiac disease [70], and red blood cell levels [71]. In patients with schizophrenia and bipolar disorder, reduced serotonin levels were seen in both disorders [72–73], further, altered circadian rhythms are believed to play a role in schizophrenia and bipolar disorder development [74]. Circadian rhythms and serotonin levels have both been discussed in relation to gut health [33–34, 75] and in patients with celiac disease, elevated levels of serotonin were described [76]. Additionally, celiac disease has been associated with mood disorders [77]. Prior studies demonstrated that serotonin-deficient mice had reduced red blood cell counts and an anaemia phenotype [78], that human red blood cells were likely influenced by circadian rhythms as they reacted to environmental stimuli [79], and lastly, patients with depression showed lower levels of red blood cell counts and higher instances of anaemia [80].

We also found evidence of two genes in our dataset within the Gene Ontology (GO) Catalogue [81–82] that have been described regarding serotonin (S8 Table). *ABCC4* is involved in platelet degranulation (GO:0002576; Reactome:R-HSA-114608), while the variant rs12209650 is intergenic between *ENSG00000276064* and *HTR1B*, where *HTR1B* has been found to enable serotonin receptor activity (GO:0004993; GO_REF:0000033) and binding (GO:0051378; GO_REF:0000107) [83]. *HTR1B* is also involved in the adenylate cyclase-inhibiting serotonin receptor signalling pathway (GO:0007198; GO_REF:0000033) [83] and negative regulation of serotonin secretion (GO:0014063) (S8 Table). Taken together, our results show that 83% of the serotonin genes we found in our results are likely adaptively introgressed and being the maximum archaic allele within their respective segment (S2 Table), highlighting the importance of serotonin and its links to circadian rhythms within modern humans. Therefore, while overt contributions to sleep and wake cycles because of archaic admixture are not found in genes linked with serotonin in our adaptive introgression results reported here, underlying mechanisms due to archaic admixture related to mood disorders and biological processes that are impacted by both circadian rhythm oscillations and serotonin levels are evident in our results. However, we also note extensive evidence of genes associated with immune function within our results, making it unclear if the association with serotonin is the driving selection event in these regions (see Gene association and function), or a combination of other processes.

### Significant GWAS associations

Two SNPs within our core haplotypes have significant GWAS associations. The *DNAAF10* core haplotype is found in the CDX at chr2:68342443-68503920 and did not display any signals of positive selection (S4 Table). The intronic variant rs6757906 has previously been linked with systolic blood pressure readings [84] (S5 Table) and is found within a gene described in the CGDB [42] as having a circadian component (S2 Table). While rs6757906 was found in frequencies over 60% in the Lahu population, it is seen predominantly in populations from the Americas, East Asia, and South Asia at frequencies over 25% (S2 Table). This marker was only seen at high frequencies in the Chagyrskaya dataset (S2 Table) and has the highest match rating to the Chagyrskaya Neanderthal within the core haplotype (S4 Table). Within the CHB population, the intronic variant rs66819621 overlaps a gene listed in the CGDB [42] and is within the same segment as the *TLR1* core haplotype (S4 Table). The variant is found in frequencies over 54% in the CHB population from the 1KGP but has worldwide allele frequencies over 15% in many 1KGP and HGDP populations (S2 Table). The archaic donor population for this region is non-specific as all three Neanderthal samples have similar match/mismatch ratios (S4 Table). This variant has been previously associated with allergic rhinitis by a study examining data from the UK Biobank [85] (S5 Table).

We identified three SNPs in our analysis that were also found at genome-wide significance in GWAS studies focused on chronotype. The intronic variant rs72799142 is found in *LINC01470* at elevated frequencies in the Brahui, Kalash, Makrani, and Sindhi populations (S2 Table). While this variant has not been reported in prior analyses specifically discussing adaptive introgression and chronotype phenotypes, the haplotypes presented in Jagoda *et al.* (2018) [86] do overlap with rs72799142. Prior GWAS analysis linked this marker with being a morning person [39] (S5 Table). The variant rs723427 is found in the *LINC01933-ENSG00000286749* intergenic region within the Surui at an allele frequency of 43.75% (S2 Table) and is also associated with being a morning person according to GWAS analysis [39] (S5 Table). McArthur *et al.* (2021) [21] previously identified a haplotype that overlapped rs723427 while Velazquez-Arcelay and colleagues (2023) [26] identified this SNP as a non-circadian variant. On the segment chr7:50426534-5089919, the intronic variant rs2190500 intersects *GRB10* in the Tu population. This SNP has also been associated with morningness [87] (S5 Table), has been highlighted previously within a gene published in the CGDB [42], and overlaps the published datasets of many studies examining archaic introgression [7, 21, 26, 86, 88]. We would also like to highlight that we found links between archaically inherited SNPs with genome-wide significant GWAS p-values with a variety of other traits (S5 Table) but are outside of the focus of this paper.

### Gene association and function

Some of the archaic variants intersecting circadian rhythm genes or showcasing circadian rhythm or chronotype traits have multiple other associated effects. For instance, the segment chr12:27534234-27967385 in the Cambodian population overlaps *MRPS35*, and the missense variant rs1127787 was discussed as being significantly linked with chronotype by Dannemann and Kelso (2017) [20] (S2 Table). However, the results from the GWAS Catalogue [43] show links with blood protein levels [89], mitochondrial DNA copy number [90–91], and type-2 diabetes [92] (S5 Table). Our results illustrate clear pleiotropy in a number of these positions (S5, S6 Tables). In this section we discuss briefly a few select association themes that are seen repeatedly throughout our annotation results (S6 Table).

Several variants in our annotation results are associated with sleep phenotypes. Variants within *ASB13*, *DAPK1*, *MIR378A*, *PPARGC1B*, and *SALL2* have all been previously tied to narcolepsy [93–94] (S6 Table). Within these, there is evidence of regional associations as the genes and trait loci, except *ASB13* found at high frequencies in Oceanic groups, are predominantly found within American populations (S3 Table). One variant, rs144380014 overlapping *PTCH1*, found only in the Melanesians, Papuans, and Naxi, (S3 Table) has associations with obstructive sleep apnea (S6 Table) in American populations [95]. Two genes, *GRB10* and *TLR1*, show evidence of generalised sleep phenotypes [96] (S6 Table), where variants associated with these genes are dispersed more generally throughout our sample populations (S3 Table). Interestingly, both genes have been associated previously with Neanderthal introgression regarding immunity-linked haplotypes [17, 21].

Links between circadian rhythm oscillations and immune function have been described previously [35–37]. In our analysis, we identified 57 genes with annotations describing immune function (S6 Table), which is over 25% of the genes found in our entire maximum allele frequency dataset (S2 Table). Six of these genes, *CCR9*, *CHST11*, *GLP1R*, *JAK1*, *KCNH7*, and *TLR1* were identified as core haplotypes, where all of them except *CHST11* and *GLP1R* had evidence of positive selection (S4 Table). All these genes have been linked with multiple traits (S6 Table) and have also been described previously in relation to archaic introgression [17, 97–103]. For example, a *CCR9* haplotype introgressed from Neanderthals conferred a susceptibility to severe COVID-19 [104], while *JAK1* has been identified with a host of other immune-response genes due to archaic admixture [105].

The connection between circadian rhythm and mental health disorders such as schizophrenia and bipolar disorder have been discussed before [74, 106]. Previous research found enrichment for schizophrenia-associated loci in Neanderthal-introgressed regions [23], however, this association has been contested when two recent studies described they found no such connection [21, 25]. Our results highlight 46 genes and two intergenic regions that have associations with schizophrenia (S6 Table). Five of these genes are found in our core haplotypes, and include *AMIGO2*, *CHST11*, *DNM1L*, *ENSG00000257643*, and *TSPAN11*, of which, only *AMIGO2* shows evidence of positive selection (S4 Table). Our maximum archaic alleles in *AMIGO2* (rs142658135), *DNM1L* (rs190280601), and *TSPAN11* (rs2241322 and rs76693329) have been associated with schizophrenia and bipolar disorder [69] (S6 Table). We found that four genes in our analysis were linked with depression (*GAD1*, *COX6CP1*, *ENSG00000275666*, and *KCNQ1*) [107–108], although none of these were core haplotypes. We also detect instances of other complex traits within our analysis, that include, but are not limited to, multiple sclerosis, Parkinson’s disease, amyotrophic lateral sclerosis (ALS/Lou Gehrig’s Disease), and Alzheimer’s (S6 Table). However, discussions regarding these are outside of the scope of this paper. Despite some contention previously regarding the archaically derived nature of some complex traits as noted above, our results show clear elevated allele frequencies at variants associated with these traits across multiple loci (S6 Table), where further research may clarify the extent of these connections.

### Limitations

Our study has several limitations. First, is that SPrime’s accuracy drops when a population has less than 15 samples for analysis [7]. This is unfortunately the case for many of the populations within the HGDP sample set. A consequence of this is that some of our windows may represent false positives. Additionally, due to the small sample size, elevated allele frequencies at many variants within these populations are seen, where other geographically similar populations with adequate sample sizes, such as in the 1KGP populations, do not show such high frequencies. We detected several introgressed variants that are either at, or nearly at, fixation (allele frequencies = 100%) in our dataset (S2 Table). For example, rs16822674 (overlapping *U3*) and rs17051049 (overlapping intergenic region *GAPDHP56-ENSG00000280059*) in the Surui are both fixed (S2 Table). However, these variants are also seen in populations with sample sizes over 15 and at allele frequencies greater than the typical introgressed archaic background frequencies [109] suggesting that the introgressed segment is correct. Further, these regions also passed our filters regarding authenticating segments and reducing instances of false positives (Materials and methods). It is possible that in some populations with small effective population sizes, such as the Surui, the effect of drift has driven archaic alleles to very high frequencies. Overall, it is important to be cautious about the interpretation of the archaic allele frequencies of some of the HGDP samples due to their very small sample sizes. A second limitation is SPrime’s masking of modern human segments found in an African reference panel [7], which has been shown to limit the detection power of archaic sequences in populations outside of the reference [88]. Therefore, we may be removing variants that may have passed our filters due to being shared with our reference population, the Yoruba in Ibadan, Nigeria (YRI). An extension of this is our filtering thresholds were quite stringent. Since we were looking for signatures of adaptive introgression, all the variants discussed here are common (40% or greater allele frequency), which means most archaic alleles will fail this filtration step. On the one hand, we can clearly exhibit instances of adaptive introgression regarding circadian rhythm and chronotype-associated variants in modern humans due to admixture with archaic hominins, on the other hand, we also removed many other very interesting segments worth exploring that may have elevated allele frequencies relative to typical archaically-introgressed levels. Future analysis should explore these limitations to help resolve some of the questions our results leave. These include utilising adequate sample sizes from more diverse human populations, exploring the effects of software on the recovery of segments with signals of adaptive introgression, investigating in detail serotonin and complex trait relationships in modern humans due to archaic introgression, deeper research into latitude clines in archaically derived regions, and lastly, examining variants with differing frequency cutoffs to see if other interesting patterns may emerge.

## Conclusions

Our paper has documented over 300 genes and intergenic segments that fall within 265 independent windows within global non-African populations. Many of these gene and intergenic segments have been described in prior studies discussing either adaptive introgression into modern humans from archaic populations or in relation to circadian rhythm and chronotype phenotypes in modern humans due to archaic introgression, confirming our results. We were able to expand on these previous analyses by investigating the extent of adaptive introgression within circadian rhythm-and chronotype-associated genomic regions within 76 worldwide populations, where previous studies have focused mostly on Eurasian populations from the 1KGP. Many of our reported genes show well documented signatures of introgression from archaic samples into modern humans, including an abundance of immunity-associated loci, complex traits including schizophrenia and bipolar disorder, and sleep associated phenotypes. Our results show clear pleiotropy, and we report in some instances the first time these regions have been described in relation with circadian rhythms and archaic introgression. Within these regions, we identified over 1,700 variants that have allele frequencies of at least 40%, and are directly matched to an archaic allele, of which, 37 genome-wide significant SNPs based on GWAS analysis were found in our dataset. Three of these GWAS variants were found to influence chronotype and the likelihood of being a morning person. In addition to these, we note that many of these significant variants were found to influence health-specific phenotypes.

We explored in detail 36 regions that we consider to be core haplotypes based on the highest allele frequency variants within these introgressed regions matching archaic alleles, having allele frequencies greater than or equal to 40%, and directly matching or falling within a region previously described as being associated with circadian rhythm or chronotype. From these, we found that 17 of these segments displayed evidence of positive selection within modern human populations, with three of these segments having evidence of positive selection in windows within 5% of the maximum archaic allele frequency variant, providing leverage to the idea that these specific regions were direct targets of adaptive introgression. We did not find definitive evidence of latitude-based clines within our core haplotype regions, instead finding clearer signals that these regions cluster more closely based on geographic similarities and previously described archaic ancestry patterns.

This study significantly advances our understanding of how archaic introgression has influenced modern human circadian rhythms and sleep patterns. By identifying a broad array of introgressed genes and intergenic segments linked to circadian functions across diverse global populations, we provide new insights into the evolutionary pressures that shaped these traits. The clear evidence of positive selection in several of these regions underscores their adaptive value. Yet, this work also suggests future, hypothesis driven work is needed to disentangle the role clinal adaptation has played in the evolution of circadian rhythms in the human lineage. This research paves the way for future studies to explore the intricate connections between circadian rhythms, mental health, and immune function, potentially leading to innovative approaches in chronomedicine and personalised healthcare.

## Materials and methods

### Modern human, Neanderthal, and Denisovan VCF files

Our modern human samples came from the previously published, phased gnomAD 1KGP + HGDP callset [41]. This unique dataset compiles the high-resolution data from the Human Genome Diversity Project (n=51 populations) and 1000 Genomes Project (n=25 populations) all mapped to GRCh38 (hg38) coordinates. Following the SPrime protocol [52], we used the YRI population from the 1KGP (n=108) as the outgroup for our analyses and combined them with each target population. We removed any non-biallelic SNPs using BCFtools v1.13 [110]. We updated all known variant IDs using the dbSNP database [111] annotation files for matching abilities in downstream analyses. Any variants with blank IDs were manually tagged with the chromosome:position:reference_alternative naming convention (i.e. chr1:15364:G_A). The archaic VCFs and their associated mask files were downloaded from the hosting sites listed in their publications. Duplicated variants were removed using PLINK2 [112].

### Introgression identification and matching to archaic sequences

To identify variants that are likely due to admixture between archaic hominins and modern humans, we used SPrime [7] with all recommended settings according to the original paper. SPrime is an archaic-reference-free software that uses a scoring parameter to identify segments in modern humans considered to be introgressed from archaic hominins. These segments are kept by the software if they are above the recommended scoring threshold of 150,000 [7]. Because the modern human genomes are mapped to hg38 coordinates and the Neanderthal and Denisovan genomes are mapped to GRCh37 (hg19), we lifted over the SPrime output files using the UCSC LiftOver Linux executable [113] to hg19. To ensure that the LiftOver [113] had not mapped any variants incorrectly, we discarded any variants that had jumped chromosomes. We used a secondary software, map_arch [52], to match our results to the archaic alleles. This software takes the SPrime output file, an archaic VCF, and the associated archaic mask file to create a new file showcasing whether the modern human variant identified by SPrime matches, mismatches, or is not comparable with the archaic genome of interest. Next, we used BCFtools [110] to generate allele frequencies for each of the modern human populations and merged these with our output using the dplyr package [114] in R [54]. We extracted the putatively introgressed segments identified in our analysis from each population, combined them together with populations from the same region according to the gnomAD sample metatable, and reduced them to the minimum number of non-overlapping segments using the GenomicRanges package v3.19 [115] in R [54].

### Chronotype and circadian rhythm datasets

We were interested in identifying if regions associated with circadian rhythm and chronotype in modern humans had any signatures of introgression from archaic hominins. Our analysis focused on compiling genes and variants linked with circadian rhythm or chronotype expression from previously reported datasets to test for these signatures. The CGDB contains over 70,000 circadian related genes identified in eukaryotic organisms [42]. We downloaded the genes found on modern human autosomes (n=1,236) from the CGDB [42] along with variants and introgressed segments published by Dannemann & Kelso (2017) [20], McArthur et al. (2021) [21], Dannemann et al. (2022) [25], and Velazquez-Arcelay et al. (2023) [26], all of which connected with circadian rhythms and chronotype expression because of archaic introgression. Additionally, we extracted all hits from the NHGRI-EBI GWAS Catalogue [43] associated with chronotype, circadian rhythm, or sleep phenotypes that had reached genome-wide significant p-values of p=5x10^-8^ or less. Since some studies will report windows suggestive of introgressed haplotypes, and others focus on just reporting variants, we opted to normalise our analysis by extracting either previously reported variants or variants found within previously described windows. Additionally, since some studies used in our analyses use different genome builds (hg19 vs. hg38), we report all our results in hg19 format to match that of the archaic samples. We extracted these variants from our results by matching the variant rsIDs using the dplyr package [114] in R [54].

### Adaptive introgression, core haplotypes, and candidates of positive selection

We were interested in identifying if any archaic variants present in the modern human genome were brought to elevated frequencies due to adaptive introgression, focusing on circadian rhythm or chronotype-associated genes. SPrime is sensitive enough to detect adaptive introgression in the human genome [7]. To test for this, we followed Browning and colleagues (2018) recommendations of selecting identified archaic segments that have 30 or more markers in the identified segment and have a match/mismatch ratio (number of matches divided by the total number of matches and mismatches) of >50% for the Neanderthals and >40% for the Denisovan [7]. After filtering out segments that failed this step, we used SnpEff [116] to annotate our VCFs and then merged this information with our population files using the rsIDs. Next, we used BEDTools [117] to intersect our population files with every other population to identify regional signatures and repeated this for each population for each archaic.

Browning and colleagues (2018) [7] identified two highly probable regions of adaptive introgression per population by first removing all variants with frequencies below 30% followed by additionally removing variants with allele frequencies 20% or more below the maximum allele frequency per segment. We applied a similar methodology, but wanted to identify possible targets of adaptive introgression in each modern human autosome that were related to circadian rhythm or chronotype. For each modern human population, we took our intersected population file and removed any putative archaic variants with allele frequencies below 40%. Next, we identified the archaic variant with the maximum archaic allele frequency per segment. Following this we then removed all variants with archaic allele frequencies more than 5% below the maximum archaic allele frequency directly upstream and downstream of that variant. We repeated this analysis for all variants that were both the maximum frequency archaic variant in their respective segment, had an allele frequency of at least 0.40, matched the archaic allele, and were either previously linked with circadian rhythm or chronotype expression or overlapped segments suggesting these signatures. This allowed us to identify what we believe are core haplotypes introgressed from archaic populations. Due to oftentimes small sample size and the effects of drift, we focused our analysis of the core haplotypes from the 1KGP populations only, along with the Papuan and Melanesian samples from the HGDP, to test for signatures of Denisovan ancestry. To give additional weight to our analysis, we also used RAiSD [55] with standard input parameters to detect evidence of positive selection within the population-specific segment containing a core haplotype. Variants determined to be in positive selection were those at the top 0.5% threshold for that chromosome, as suggested by the RAiSD documentation [55].

### Archaic donor populations

The ratio of the number of matches and mismatches can be compared to identify whether introgressed segments are from Neanderthals or Denisovans [7, 52]. We tested for this by taking segments with 30 or more variants where segments that are believed to be of Neanderthal affinity will have match/mismatch ratios greater than 60% to a Neanderthal sample coinciding with a match/mismatch ratio below 40% for the Denisovan [7]. Similarly, segments likely of Denisovan origin will have a match/mismatch ratio more than 40% to the Denisovan and below 30% with the Neanderthals [7]. We first excluded segments that did not pass our adaptive introgression thresholds for each population relative to each archaic sample, and then identified which of these segments are clearly introgressed from one archaic sample relative to the others. When a segment passed the donor thresholds and had a segment match/mismatch ratio more than 5% higher relative to the other three archaic samples, we inferred that the putative archaic donor is closest to that archaic population. If the match/mismatch ratio is within 5% relative to the other archaic samples, we consider that to be inconclusive and is of archaic affinity generally. We applied these calculations to our core haplotypes. To visualise these affinities, we generated contour plots based on the scripts provided by Zhou and Browning (2021) [52] in R [54] using the kde2d function from MASS [58] for core haplotypes with evidence of positive selection.

### Gene and Variant Phenotype Associations

For all genes described in our results we attempted to attribute a phenotypic association. Variant analysis was done using the GRCh37 search in SNPnexus [59–60] and in VEP for the GRCh37 Release 112 [61]. Gene ontology information was compiled from the GO Consortium data [81–82] on genes downloaded from BioMart in the GRCh37 Release 112 [61].

## Supporting information

**S1 Text. Supporting information and methodology.** A description of the number of recovered variants in our pipeline from previously published results reporting adaptive introgression in modern humans due to archaic admixture. Methodological outlines for our haplotype networks and ancestral recombination graphs are also included. (DOCX)

**S1 Table. Introgressed segments.** Segments represent putatively introgressed regions identified by SPrime [7] that have been merged into non-overlapping windows between geographically similar populations. Geographic regions were defined by the gnomAD metatable. Coordinates are sorted in genomic order and in hg19 format. (XLSX)

**S2 Table. Maximum archaic frequency variants per segment.** Archaic segments that contain variants where any of the following are true: matches a previously reported circadian rhythm variant, is found within previously identified regions in other prior studies discussing archaic introgression and circadian rhythm, or overlaps a region from the CGDB database. Variants in these segments are the maximum archaic allele frequency within their segments, have >=40% allele frequency within at least one population, and directly match the archaic allele. The segments are sorted in chromosomal order and include intersection and gene annotation results. Each archaic sample has its own tab for simplicity due to the size of the table and to show differences in the introgressing segments. Coordinates are in hg19 format. (XLSX)

**S3 Table. High frequency archaic variants.** Archaic segments that contain variants where any of the following are true: matches a previously reported archaic-circadian rhythm variant, is found within previously identified regions in other prior studies discussing archaic introgression, and circadian rhythm, or overlaps a region from the CGDB database. Variants in these segments are also >=40% allele frequency within at least one population and directly match the archaic allele. Note that these are different from S2 Table in that these variants not necessarily the maximum archaic allele in their segments. The segments are sorted in chromosomal order and include intersection and gene annotation results. Each archaic sample has its own tab for simplicity due to the size of the table and to show differences in the introgressing segments. Coordinates are in hg19 format. (XLSX)

**S4 Table. List of core haplotypes.** The core haplotypes were generated by identifying variants that had archaic allele frequencies >=40%, were the maximum archaic allele frequency variant in their segment and had associations with circadian rhythm and/or chronotype. We generated windows around the main variant with a 5% allele frequency threshold directly upstream and downstream of the variant to create the core. Evidence of positive selection was generated using RAiSD [55]. Blue highlighting shows positive selection was detected within the segment identified by SPrime [7] but outside of the core, while green shows positive selection directly within the core. Coordinates are in hg19 format. (XLSX)

**S5 Table. Genome-wide significant variants within GWAS analyses.** Archaic segments containing significant GWAS associated variants where the archaic allele is >=40% allele frequency in at least one population. The segments are sorted in chromosomal order and coordinates are in hg19 format. GWAS data obtained from NHGRI-EBI GWAS Catalogue [43]. (XLSX)

**S6 Table. Variant annotations.** Variant annotation results combined from SNPnexus [59–60] and VEP [61]. The table is sorted in alphabetical order based on gene name and the coordinates are in hg19 format. PubMed ID (or reference) to the study is provided where possible based on the downloaded data, a “0” indicates no reference was given from either source. (XLSX)

**S7 Table. Latitude cline.** Examination of latitudinal cline within core haplotypes. Population-specific regions, and associated maximum archaic allele frequencies, were extracted from each population based on the whole segment listed in S3 Table. Latitudes were obtained from the gnomAD sample metatable. Not all populations have putatively introgressed segments overlapping core haplotypes, as these populations have segments that match the African outgroup alleles (the YRI). These are denoted with a 0 for their allele frequencies. The allele frequencies are taken from the samples before filtering for authentic segments (see Materials and methods). Correlation statistics were done using PAST [118] and are reported using absolute latitude, where negatives were removed and input as a positive number. (XLSX)

**S8 Table. Gene ontology (GO) data.** GO data for genes within our results downloaded from BioMart [61]. The table is in alphabetical order based on the gene name. Missing data based on the downloaded information will contain a “-” in the column. (XLSX)

**S1 Fig. *AMIGO2* linear regression graph.** Comparison of maximum archaic allele frequency (x-axis) against absolute latitude (y-axis) for the *AMIGO2* core haplotype. There was no significant relationship (p=0.13896) with a low coefficient of determination (r^2^ = 0.029348). The line of best fit is sloped negatively. The plot and summary statistics were generated using PAST [118]. (TIF)

**S2 Fig. *CCR9* linear regression graph.** Comparison of maximum archaic allele frequency (x-axis) against absolute latitude (y-axis) for the *CCR9* core haplotype. There was no significant relationship (p=0.10945) with a low coefficient of determination (r^2^ = 0.034256). The line of best fit is sloped positively. The plot and summary statistics were generated using PAST [118]. (TIF)

**S3 Fig. *CEACAM1-LIPE-AS1* linear regression graph.** Comparison of maximum archaic allele frequency (x-axis) against absolute latitude (y-axis) for the *CEACAM1-LIPE-AS1* core haplotype. There was a significant relationship (p=0.028042) with a low coefficient of determination (r^2^ = 0.063541). The line of best fit is sloped negatively. The plot and summary statistics were generated using PAST [118]. (TIF)

**S4 Fig. *ENSG00000286749* linear regression graph.** Comparison of maximum archaic allele frequency (x-axis) against absolute latitude (y-axis) for the *ENSG00000286749* core haplotype. There was no significant relationship (p=0.29001) with a low coefficient of determination (r^2^ = 0.015117). The line of best is sloped positively. The plot and summary statistics were generated using PAST [118]. (TIF)

**S5 Fig. *ENSG00000279193-ENSG00000276122* linear regression graph.** Comparison of maximum archaic allele frequency (x-axis) against absolute latitude (y-axis) for the *ENSG00000279193-ENSG00000276122* core haplotype. There was no significant relationship (p=0.16315) with a low coefficient of determination (r^2^ = 0.026111). The line of best is sloped negatively. The plot and summary statistics were generated using PAST [118]. (TIF)

**S6 Fig. *JAK1* linear regression graph.** Comparison of maximum archaic allele frequency (x-axis) against absolute latitude (y-axis) for the *JAK1* core haplotype. There was no significant relationship (p=0.06091) with a low coefficient of determination (r^2^ = 0.046662). The line of best is sloped negatively. The plot and summary statistics were generated using PAST [118]. (TIF)

**S7 Fig. *KCNH7* linear regression graph.** Comparison of maximum archaic allele frequency (x-axis) against absolute latitude (y-axis) for the *KCNH7* core haplotype. There was no significant relationship (p=0.31644) with a low coefficient of determination (r^2^ = 0.013561). The line of best is sloped negatively. The plot and summary statistics were generated using PAST [118]. (TIF)

**S8 Fig. *LINC01107-LINC01937* linear regression graph.** Comparison of maximum archaic allele frequency (x-axis) against absolute latitude (y-axis) for the *LINC01107-LINC01937* core haplotype. There was a significant relationship (p=0.035543) with a low coefficient of determination (r^2^ = 0.058346). The line of best is sloped positively. The plot and summary statistics were generated using PAST [118]. (TIF)

**S9 Fig. *MIER3* linear regression graph.** Comparison of maximum archaic allele frequency (x-axis) against absolute latitude (y-axis) for the *MIER3* core haplotype. There was no significant relationship (p=0.075055) with a low coefficient of determination (r^2^ = 0.042196). The line of best is sloped negatively. The plot and summary statistics were generated using PAST [118]. (TIF)

**S10 Fig. *RN7SL423P-ENSG00000232337* linear regression graph.** Comparison of maximum archaic allele frequency (x-axis) against absolute latitude (y-axis) for the *RN7SL423P-RNSG00000232337* core haplotype. There was no significant relationship (p=0.098013) with a low coefficient of determination (r^2^ = 0.036559). The line of best is sloped positively. The plot and summary statistics were generated using PAST [118]. (TIF)

**S11 Fig. *ROR2* linear regression graph.** Comparison of maximum archaic allele frequency (x-axis) against absolute latitude (y-axis) for the *ROR2* core haplotype. There was a significant relationship (p=0.022986) with a low coefficient of determination (r^2^ = 0.067914). The line of best is sloped positively. The plot and summary statistics were generated using PAST [118]. (TIF)

**S12 Fig. *RPSAP11-ENSG00000261572* linear regression graph.** Comparison of maximum archaic allele frequency (x-axis) against absolute latitude (y-axis) for the *RPSAP11-ENSG00000261572* core haplotype. There was no significant relationship (p=0.84219) with a low coefficient of determination (r^2^ = 0.0005391). The line of best is sloped negatively. The plot and summary statistics were generated using PAST [118]. (TIF)

**S13 Fig. *SUSD1* linear regression graph.** Comparison of maximum archaic allele frequency (x-axis) against absolute latitude (y-axis) for the *SUSD1* core haplotype. There was no significant relationship (p=0.4878) with a low coefficient of determination (r^2^ = 0.00652766). The line of best is sloped negatively. The plot and summary statistics were generated using PAST [118]. (TIF)

**S14 Fig. *TIAM2* linear regression graph.** Comparison of maximum archaic allele frequency (x-axis) against absolute latitude (y-axis) for the *TIAM2* core haplotype. There was no significant relationship (p=0.59898) with a low coefficient of determination (r^2^ = 0.0037553). The line of best is sloped negatively. The plot and summary statistics were generated using PAST [118]. (TIF)

**S15 Fig. *TLR1* linear regression graph.** Comparison of maximum archaic allele frequency (x-axis) against absolute latitude (y-axis) for the *TLR1* core haplotype. There was a significant relationship (p=0.00000758) with a low coefficient of determination (r^2^ = 0.23862). The line of best is sloped positively. The plot and summary statistics were generated using PAST [118]. (TIF)

**S16 Fig. *AMIGO2* maximum archaic allele frequency map.** The maximum archaic allele frequency of each population in our analysis for *AMIGO2* plotted using rnaturalearth [65], sf [66–67], and ggplot2 [53] in R [54].

**S17 Fig. *CCR9* maximum archaic allele frequency map.** The maximum archaic allele frequency of each population in our analysis for *CCR9* plotted using rnaturalearth [65], sf [66–67], and ggplot2 [53] in R [54].

**S18 Fig. *CEACAM1-LIPE-AS1* maximum archaic allele frequency map.** The maximum archaic allele frequency of each population in our analysis for *CEACAM1-LIPE-* plotted using rnaturalearth [65], sf [66–67], and ggplot2 [53] in R [54].

**S19 Fig. *ENSG00000286749* maximum archaic allele frequency map.** The maximum archaic allele frequency of each population in our analysis for *ENSG00000286749* plotted using rnaturalearth [65], sf [66–67], and ggplot2 [53] in R [54].

**S20 Fig. *ENSG00000279193-ENSG00000276122* maximum archaic allele frequency map.** The maximum archaic allele frequency of each population in our analysis for *ENSG00000279192-ENSG00000276122* plotted using rnaturalearth [65], sf [66–67], and ggplot2 [53] in R [54].

**S21 Fig. *JAK1* maximum archaic allele frequency map.** The maximum archaic allele frequency of each population in our analysis for *JAK1* plotted using rnaturalearth [65], sf [66–67], and ggplot2 [53] in R [54].

**S22 Fig. *KCNH7* maximum archaic allele frequency map.** The maximum archaic allele frequency of each population in our analysis for *KCNH7* plotted using rnaturalearth [65], sf [66–67], and ggplot2 [53] in R [54].

**S23 Fig. *LINC01107-LINC01937* maximum archaic allele frequency map.** The maximum archaic allele frequency of each population in our analysis for *LINC01107-LINC01937* plotted using rnaturalearth [65], sf [66–67], and ggplot2 [53] in R [54].

**S24 Fig. *MIER3* maximum archaic allele frequency map.** The maximum archaic allele frequency of each population in our analysis for *MIER3* plotted using rnaturalearth [65], sf [66–67], and ggplot2 [53] in R [54].

**S25 Fig. *RN7SL432P-ENSG00000232337* maximum archaic allele frequency map.** The maximum archaic allele frequency of each population in our analysis for *RN7SL432P-ENSG00000232337* plotted using rnaturalearth [65], sf [66–67], and ggplot2 [53] in R [54].

**S26 Fig. *ROR2* maximum archaic allele frequency map.** The maximum archaic allele frequency of each population in our analysis for *ROR2* plotted using rnaturalearth [65], sf [66–67], and ggplot2 [53] in R [54].

**S27 Fig. *RPSAP11-ENSG00000261572* maximum archaic allele frequency map.** The maximum archaic allele frequency of each population in our analysis for *RPSAP11-ENSG00000261572* plotted using rnaturalearth [65], sf [66–67], and ggplot2 [53] in R [54].

**S28 Fig. *SUSD1* maximum archaic allele frequency map.** The maximum archaic allele frequency of each population in our analysis for *SUSD1* plotted using rnaturalearth [65], sf [66–67], and ggplot2 [53] in R [54].

**S29 Fig. *TIAM2* maximum archaic allele frequency map.** The maximum archaic allele frequency of each population in our analysis for *TIAM2* plotted using rnaturalearth [65], sf [66–67], and ggplot2 [53] in R [54].

**S30 Fig. *TLR1* maximum archaic allele frequency map.** The maximum archaic allele frequency of each population in our analysis for *TLR1* plotted using rnaturalearth [65], sf [66–67], and ggplot2 [53] in R [54].

**S31 Fig. *CEACAM1-LIPE-AS1* global ancestral recombination graph.** Ancestral recombination graph for rs184528844 inside of *CEACAM1-LIPE-AS1* for the Oceanic populations plotted against all non-African populations from the 1KGP with the YRI as a proxy for sub-Saharan African populations. The plot was generated using Relate [57]. (TIF)

**S32 Fig. *CEACAM1-LIPE-AS1* regional ancestral recombination graph.** Ancestral recombination graph for rs184528844 inside of *CEACAM1-LIPE-AS1* for the Oceanic populations plotted against just the YRI. The plot was generated using Relate [57]. (TIF)

**S33 Fig. *CCR9* global ancestral recombination graph.** Ancestral recombination graph for rs71327015 inside of *CCR9* for the South Asian populations plotted against all non-African populations from the 1KGP with the YRI as a proxy for sub-Saharan African populations. The plot was generated using Relate [57]. (TIF)

**S34 Fig. *CCR9* regional ancestral recombination graph.** Ancestral recombination graph for rs71327015 inside of *CCR9* for the South Asian populations plotted against just the YRI. The plot was generated using Relate [57]. (TIF)

**S35 Fig. *JAK1* global ancestral recombination graph.** Ancestral recombination graph for rs377425962 inside of *JAK1* for the Oceanic populations plotted against all non-African populations from the 1KGP with the YRI as a proxy for sub-Saharan African populations. The plot was generated using Relate [57]. (TIF)

**S36 Fig. *JAK1* regional ancestral recombination graph.** Ancestral recombination graph for rs377425962 inside of *JAK1* for the South Asian populations plotted against just the YRI. The plot was generated using Relate [57]. (TIF)

